# Early patterns of functional brain development associated with autism spectrum disorder in tuberous sclerosis complex

**DOI:** 10.1101/578831

**Authors:** Abigail Dickinson, Kandice J. Varcin, Mustafa Sahin, Charles A. Nelson, Shafali S. Jeste

**Affiliations:** UCLA Semel Institute of Neuroscience and Human Behavior, David Geffen School of Medicine, 760 Westwood Plaza, Los Angeles, CA, 90095; Telethon Kids Institute, University of Western Australia, 100 Roberts Road, Subiaco, WA, 6008, Australia; Department of Neurology, Boston Children’s Hospital, Translational Neuroscience Center, 300 Longwood Avenue, Boston, MA, 02115; Division of Developmental Medicine, Boston Children’s Hospital, Harvard Medical School & Harvard Graduate School of Education, Harvard University, 1 Autumn Street, Boston, MA, 02215

**Keywords:** Tuberous sclerosis complex, autism spectrum disorder, functional connectivity, infancy, electroencephalography, cognitive function, alpha oscillations

## Abstract

Around half of infants with tuberous sclerosis complex (TSC) develop autism. Here, using EEG, we find that there is a reduction in communication between brain regions during infancy in TSC, and that the infants who show the largest reductions are those who later develop autism. Being able to identify infants who show early signs of disrupted brain development may improve the timing of early prediction and interventions in TSC, and also help us to understand how early brain changes lead to autism.

**Abstract:** Tuberous sclerosis complex (TSC) is a rare genetic disorder that confers a high risk for autism spectrum disorders (ASD), with behavioral predictors of ASD emerging early in life. Deviations in structural and functional neuronal connectivity are highly implicated in both TSC and ASD.

For the first time, we explore whether electroencephalographic (EEG) measures of network function precede or predict the emergence of ASD in TSC. We determine whether altered brain function (1) is present in infancy in TSC, (2) differentiates infants with TSC based on ASD diagnostic status, and (3) is associated with later cognitive function.

We studied 35 infants with TSC (N=35), and a group of typically developing infants (n=20) at 12 and 24 months of age. Infants with TSC were later subdivided into ASD and non-ASD groups based on clinical evaluation. We measured features of spontaneous alpha oscillations (6-12Hz) that are closely associated with neural network development: alpha power, alpha phase coherence (APC) and peak alpha frequency (PAF).

Infants with TSC demonstrated reduced interhemispheric APC compared to controls at 12 months of age, and these differences were found to be most pronounced at 24 months in the infants who later developed ASD. Across all infants, PAF at 24 months was associated with verbal and non-verbal cognition at 36 months.

Associations between early network function and later neurodevelopmental and cognitive outcomes highlight the potential utility of early scalable EEG markers to identify infants with TSC requiring additional targeted intervention initiated very early in life.

## Introduction

Tuberous sclerosis complex (TSC) is a rare autosomal-dominant genetic syndrome caused by the inactivation of TSC1 or TSC2 genes. The TSC1/TSC2 protein complex plays a critical role in regulating cell growth and proliferation via mechanistic target of rapamycin (mTOR) signaling. In TSC, MTOR remains constitutively active, resulting in unrestricted cell growth and proliferation throughout the body. While TSC is considered a multisystem disorder (affecting the kidneys, heart, eyes, lungs, and skin), the central nervous system is consistently and prominently affected (Curatolo, Bombardieri, & Jozwiak, 2008). The majority of individuals with TSC have hamartomas in the brain, including cortical tubers, subependymal nodules and/or subependymal giant-cell astrocytomas (Northrup, Krueger, & International Tuberous Sclerosis Complex Consensus Group, 2013; Roth et al., 2013), with prevalence estimates of epilepsy up to 90% (Chu-Shore, Major, Camposano, Muzykewicz, & Thiele, 2010; Devlin, Shepherd, Crawford, & Morrison, 2006). Neurodevelopmental or neuropsychiatric disorders (referred to as TSC-associated neuropsychiatric disorders [TAND]; (de Vries et al., 2015; Leclezio, Jansen, Whittemore, & de Vries, 2015)) are common. 25-60% of children with TSC will receive a diagnosis of Autism Spectrum Disorder (ASD) (Capal et al., 2017; Granader et al., 2010; Jeste, Sahin, Bolton, Ploubidis, & Humphrey, 2008; Vignoli et al., 2015), and more than 50% have some degree of cognitive impairment (Curatolo, Moavero, & de Vries, 2015; Joinson et al., 2003).

TSC is often diagnosed in utero or shortly after birth through identification of either cardiac or cortical hamartomas (Datta, Hahn, & Sahin, 2008; Davis et al., 2017), facilitating the examination of these infants well before diagnoses of ASD or intellectual disability (ID) are made. Longitudinal studies have revealed early behavioral differences between infants and toddlers with TSC who later receive an ASD diagnosis and those who do not (Jeste et al., 2016; McDonald et al., 2017). Infants who develop ASD are found to demonstrate social communication deficits by 9 months of age (McDonald et al., 2017) and lower cognitive function by 12 months of age. Interestingly, toddlers not diagnosed with ASD demonstrate social communication profiles indistinguishable from typically developing (TD) children, despite having comparable cortical tuber burden and epilepsy severity to those with ASD (Jeste et al., 2014). Given the high risk of neurodevelopmental disorders in TSC, with symptoms emerging in infancy, we asked whether there are neurobiological markers of atypical brain development that relate to and predict neurodevelopmental disorders in TSC.

Commensurate with our understanding of the neurobiological mechanisms underlying neurodevelopmental disorders as a whole (Geschwind & Levitt, 2007; Jeste, Frohlich, & Loo, 2015), it is most likely that altered patterns of neuronal connectivity (outside of cortical tubers, subependymal nodules and astrocytomas) account for atypical neurodevelopment in TSC (Ebrahimi-Fakhari & Sahin, 2015; Jülich & Sahin, 2014; Peters et al., 2012). The critical role of TSC1/2 proteins in neuronal connectivity is supported by evidence from preclinical studies that demonstrate abnormalities in neuronal migration and morphology (Knox et al., 2007; Nie et al., 2010), synaptic plasticity and function (Ehninger et al., 2008; Zeng et al., 2007), excitation-inhibition balance (Bateup et al., 2013), and myelination patterns (Ercan et al., 2017) in the context of a TSC mutation. In addition, structural imaging studies in individuals with TSC demonstrate widespread alterations in structural connections (Baumer et al., 2015; Krishnan et al., 2010; Lewis et al., 2012; Peters et al., 2012, 2012), relevant to the variability seen in neurodevelopmental outcomes. Using an unbiased whole-brain connectivity analysis, Im et al. (2016) identified altered structural connectivity in TSC with markedly reduced inter-hemispheric connectivity (aged 3-24 years) compared to a control sample, most prominent in those with developmental delay (Im et al., 2015). Similarly, deficits in the integrity of white matter microstructure are found to be more pronounced in individuals with TSC who also have an ASD diagnosis (Baumer et al., 2017; Lewis et al., 2012; Peters et al., 2012). Finally, and highly relevant to the present study, a pattern of functional hypo-connectivity is identified using electroencephalography (EEG) in individuals with TSC across a wide age range, with hypoconnectivity (in some cortical areas) more prominent in individuals with both TSC and ASD (Peters et al., 2013)..

EEG offers a scalable and feasible method to capture the specific changes in brain dynamics that occur during infancy. EEG measures brain function in three important dimensions: space, time and frequency (Kida, Tanaka, & Kakigi, 2016). Here we focus on oscillations which occur within the alpha range (6-12Hz), which represent the dominant signal in task-independent environments (Guggisberg et al., 2008), and have been well-characterized during early infancy (Saby & Marshall, 2012). In addition to measuring the power of alpha oscillations, we focus on two measures of alpha activity, alpha phase coherence (APC) and peak alpha frequency (PAF), that are specifically sensitive to changes in neuronal connectivity that occur over the course of typical brain maturation, and in clinical populations. Across the oscillatory signals that co-ordinate large scale neural communication, the coherence of neural signals in the alpha range, APC, is demonstrated to be the most sensitive to both the healthy development of (Smit et al., 2012), and disruptions to (Hinkley et al., 2011), functional and structural connectivity. Similarly, the frequency at which maximal power occurs within the alpha band, PAF, shows well-characterized increases with chronological age in TD children (Chiang, Rennie, Robinson, van Albada, & Kerr, 2011; Cragg et al., 2011; Dustman, Shearer, & Emmerson, 1999; Miskovic et al., 2015; Somsen, van’t Klooster, van der Molen, van Leeuwen, & Licht, 1997; Stroganova, Orekhova, & Posikera, 1999). In ASD, PAF relates to cognitive function, not age (Dickinson, DiStefano, Senturk, & Jeste, 2017). In addition to age and cognition, both APC and PAF are associated with white matter architecture (axon growth and myelination) (Jann et al., 2012; Teipel et al., 2009), further suggesting that these metrics index the development of neural networks (Wolfgang Klimesch, Sauseng, & Hanslmayr, 2007; Valdés-Hernández et al., 2010).

As part of a longitudinal study of TSC, we utilized high-density EEG measures of spontaneous alpha oscillations to characterize the development of functional connectivity across first two years of life and to determine whether these early biomarkers informed neurodevelopmental outcomes. More specifically, we asked: 1) Do alpha oscillations (power, PAF and PAC) differ between infants with and without TSC?; (2) Do alpha oscillations (power, PAF and PAC) differentiate infants with TSC who develop ASD from those that do not?; and 3) Does PAF measured during infancy predict later cognitive function? Data from this study could reveal the earliest manifestations of atypical brain development that may precede and predict the development of ASD and ID in TSC. Such investigations can help elucidate mechanisms underlying the distinct (and often diverse) neurodevelopmental outcomes in TSC, identify early predictive biomarkers of neurodevelopmental disorders, further stratify infants by risk for neurodevelopmental disorders, and provide objective measures of change with early intervention in high-risk infants.

## Method

### Participants

Infants were enrolled in a longitudinal, two-site study examining early development in TSC (Jeste et al., 2016, 2014; McDonald et al., 2017). Infants with confirmed TSC diagnoses (Northrup et al., 2013) were recruited through specialty clinics, newborn nurseries, pediatricians’ offices and the Tuberous Sclerosis Alliance. TD control infants (described below) were recruited through institutional review board-approved databases.

Infants visited one of the two study sites throughout the first two years of life (at 6, 9, 12 and/or 24 months of age) to undergo EEG recording. At 18, 24 and/or 36 months, infants underwent behavioral assessments to determine ASD symptoms (described in detail below), which was followed by a Best Clinical Estimate diagnostic process. A schematic diagram of the study protocol can be found in the supplementary materials. Although we collected EEG data in a small subset of infants at age 6 and 9 months, in order to achieve adequate sample sizes to interrogate statistical differences between groups, we focused on 12 and 24 month time points.

To be included in the present analyses, participants had to have EEG data for at least one of the two time points (12 and 24 months). Exclusion criteria for TD infants included prematurity (<37 weeks gestational age), birth trauma, developmental concerns, or a first-degree relative with ASD or ID. It should be noted that while ‘typically developing’ may not traditionally be used to describe infants, our later follow-up time points were used to confirm that no developmental concerns were present in the TD group at 36 months of age. The study received ethical approval from both study sites. Parents provided informed written consent, in accordance with the Declaration of Helsinki.

TD (N=22) and TSC (N=36) infants completed an EEG recording session at 12 or 24 months. Of this sample, 2 TD participants and 1 TSC participant were excluded from the current analyses due to extensive EEG artifact. Final groups therefore consisted of 35 TSC (12 months: n=18; 24 months: n=28), and 20 TD infants (12 months: n=20; 24 months: n=12). Clinical genetics reports were available for n=21 (60%) of the total TSC sample, with all TSC infants meeting clinical criteria for TSC (Northrup et al., 2013). Demographic and developmental information for the sample is summarized in Table 1.

**Table 1.** Demographic participant information.

|  | **TD Control** | **TSC** | **ASD-** | **ASD+** |
| --- | --- | --- | --- | --- |
|  | **(n=20)** | **(n=35)** | **(n=14)** | **(n=17)** |
| **Sex** |  |  |  |  |
| Number female (% female). | 11 f (55%) | 9 f (26%) | 4 f (29%) | 4 f (24%) |
| **12 Month EEG** |  |  |  |  |
| Number completed (percentage completed). | n=20 (100%) | n=18 (51%) | N=7 (50%) | n=8 (47%) |
| **24 Month EEG** |  |  |  |  |
| Number completed (percentage completed). | n=12 (60%) | n=28 (80%) | n=13 (93%) | n=14 (82%) |
| **36 Months Verbal DQ** | M=115.20 | M=68.92 | M=82.74 | M=47.42 |
| Mean (SD); Number completed (percentage completed) | (SD=30.68)<br>n=10 (50%) | (SD=30.38)<br>n=23 (66%) | (SD=22.57)<br>n=14 (100%) | (SD=29.25)<br>n=9 (53%) |
| **36 Months Non-Verbal DQ** | M=113.87 | M=73.32 | M=84.40 | M=56.09 |
| Mean (SD); Number completed (percentage completed) | (SD=24.34)<br>n=10 (50%) | (SD=24.27)<br>n=23 (66%) | (SD=20.78)<br>n=14 (100%) | (SD=19.24)<br>n=9 (53%) |
| **ASD Outcome** |  |  |  |  |
| Mean Calibrated severity score (SD); Number completed (percentage completed) | M=1.05<br>(SD=0.52)<br>n=19 (95%) | M=4.3<br>(SD=2.77)<br>n=30 (86%) | M=1.71<br>(SD=.82)<br>n=14 (100%) | M=6.24<br>(SD=2.05)<br>n=17 (100%) |

It should be noted that males were over-represented in the TSC group. While it has been shown that many behavioral features (including the presence of ASD and ID) are distributed equally across both sexes in TSC (Curatolo, Porfirio, Manzi, & Seri, 2004; de Vries, Hunt, & Bolton, 2007), larger samples in the future (described in the discussion) will allow the exploration of sex-related differences in early brain function in TSC.

#### Seizures, Tubers & Medication

Thirty-two (91%) of participants with TSC had seizure data available through parent-report and/or medical records. Of these individuals, the majority (29/32; 91%) had seizures. Age of onset ranged from day of birth to 22 months, with a median age of onset at 4 months (interquartile range = 4.56 months). Of those with seizures, 62.5% (20/32) had infantile spasms. Information regarding the presence of cortical tubers was present for 80% of the TSC sample (n=28). Of the participants with available tuber information, 28 (100%) had tubers evident on clinical MRI. Of infants with available medication information (26/32; 81%), 92% (24/26) had taken antiepileptic medications (AEDs) at either the 9, 12, and/or 24 month assessments. Only the 2 infants with TSC who had no history of seizures were naïve to AEDs. No history of seizures or medication use were reported for any TD infant. See supplementary tables S1 and S2 for additional details regarding seizure history and medication.

### Behavioral assessments

#### Autism symptoms and diagnosis

Clinical ASD symptoms were assessed using Modules 1 and 2 of the Autism Diagnostic Observation Schedule (ADOS; (Lord et al., 2000). Calibrated severity scores (CSS) were calculated in order to facilitate comparison of scores across modules (Gotham, Risi, Pickles, & Lord, 2007). A diagnostic outcome of ASD (ASD+) was based on both meeting the ADOS cut-off score for ASD (CSS ≥ 4), and a clinical best estimate judgement carried out by a trained clinician. Infants with TSC who did not show elevated ASD symptoms are described as ASD−.

Nineteen (95%) TD infants underwent an ADOS assessment. These were carried out at either 18 (n=3; 15.8%), 24 (n=6; 31.6%) or 36 months (n=10; 52.6%). No TD infants met criteria for ASD. Of the 36 TSC infants included in the present study, 4 (11%) did not complete an ADOS assessment. These infants were included in the main group comparison (TSC compared to TD infants) but not included in any sub-analyses of ASD+ compared to ASD−. Of the four children with TSC who did not complete an ADOS assessment, one could not have the ADOS administered in their native language, and three were lost to follow-up. Of the 31 TSC infants that received an ADOS assessment at either 18 (n=1; 3%), 24 (n=6; 19%) or 36 months (n=24; 77%), 17 were deemed to have met ASD criteria based on Best Clinical Estimate (55%), and 14 did not (45%). This is consistent with previously reported prevalence rates of ASD in TSC (Jeste et al., 2008).

#### Cognitive Function

Cognitive function was assessed at 36 months of age using a standardized measure of cognitive and motor development, the Mullen Scales of Early Learning (MSEL; (Mullen, 1995). A verbal developmental quotient (VDQ) was calculated using the average age equivalent score of the Receptive Language and Expressive Language subscale scores (divided by chronological age), and a non-verbal developmental quotient (NVDQ) was calculated using the average age equivalent score of the Visual Reception and Fine Motor subscale scores (divided by chronological age) (Akshoomoff, 2006). Cognitive function measures were only available for 50% of the TD group and 66% of the TSC group, due to a large number of participants being lost to follow up before cognitive function was assessed at 36 months (see Table 1).

### EEG Acquisition and Processing

#### EEG Recording

Continuous EEG data were recorded using a high density 128-channel HydroCel Geodesic Sensor Net (Electrical Geodesics Inc., Eugene, OR). EEG was recorded for five minutes in a dark, sound-attenuated room while infants sat in their caregiver’s lap. Consistent with many other studies in developmental populations, a research assistant aimed to keep the infant settled and content for the duration of the recording by blowing bubbles (Dawson et al. 1995; McEvoy et al. 2015; Stroganova et al. 1999; Tierney et al. 2012; Webb et al. 2015; Gabard-Durnam, 2015; Levin, 2017). Net Station 4.4.5 software was used to record from a Net Amps 300 amplifier with a low-pass analog filter cutoff frequency of 6 KHz. Data were sampled at 500Hz and referenced to vertex at the time of recording. Electrode impedances were kept below 100 KΩ.

#### EEG Data Pre-processing

All offline data processing and analyses were performed using EEGLAB (Delorme & Makeig, 2004), and in-house MATLAB scripts. Data were high pass filtered to remove frequencies below 1Hz and low pass filtered to remove frequencies above 50Hz, using a finite impulse response filter implemented in EEGLAB. Continuous data were then visually inspected, and noisy channels were removed. Following channel removal, data were interpolated to the international 10-20 system 25 channel montage (Jasper, 1958). Sections of data that showed electromyogram (EMG) or other non-stereotyped artifacts, based on visual inspection of the signal, were then removed from the recording. Independent component analysis (ICA), a statistical blind source separation technique (Makeig, Jung, Bell, Ghahremani, & Sejnowski, 1997), was implemented to remove electrooculogram (EOG) and other stereotyped artifacts from the data. After decomposing the data into maximally independent components (IC), the scalp topography and time course of each IC was visually inspected. Any IC that represented a non-neural source (including EMG, EOG and line noise) was removed from the data. The experimenter was blinded to all participant details throughout the data cleaning process.

Data were then separated into three-second epochs. The minimum amount of artifact free data available across participants was 46 seconds, thus the first 45 seconds (15 epochs) of data from every participant underwent analyses. This data length represents an appropriate minimum threshold to gain reliable estimates of the characteristics of spontaneous EEG (Dickinson et al., 2017; Gudmundsson, Runarsson, Sigurdsson, Eiriksdottir, & Johnsen, 2007).

#### EEG data processing: Alpha Power

Welch’s method, implemented using 2-second Hamming windows with 50% overlap, was used to compute spectral power from cleaned EEG for each of the 25 electrodes. The resulting power spectra had approximately 0.5 Hz frequency resolution. Relative power was calculated for each region of interest by determining the proportion of total spectral power (1–50 Hz) accounted for by each frequency bin. Relative power values were then summed for frequencies within the alpha (6-12Hz) range.

#### EEG data Processing: Peak Alpha Frequency (PAF)

PAF was computed for three regions of interest using the computed power spectra (described above) from six channels (F3, F4, C3, C4, O1 and O2), with each region defined by averaging the power spectra from two channels (i.e., F3, F4 = frontal; C3, C4 = central; O1, O2 = occipital). A robust curve fitting procedure (fully described in Dickinson et al., 2017) was used to quantify peak alpha frequency in an objective manner that is free of bias towards lower frequencies (Neto, Allen, Aurlien, Nordby, & Eichele, 2015). If the curve-fitting procedure failed, data were omitted from further analysis. Chi-square analyses revealed that there were no group differences in peak presence between TSC and TD groups, with 9.63% of participants at 12 months, and 3.3% of participants at 24 months having at least one region omitted from analysis.

#### EEG data processing: Alpha Phase Coherence (APC)

Cleaned EEG data underwent surface Laplacian transform using realistic head geometry, based on the average head radius of participants in the present sample (12 months = 7.3cm; 24 months = 7.8cm). The surface Laplacian is the second spatial derivative of the scalp recorded potentials for each electrode, thus transforming the scalp-recorded EEG into estimates of current source density (CSD). We employed a spline flexibility constant of m=3, which represents adequate flexibility to prevent distortion of the original data (Kayser & Tenke, 2015).

Coherence analyses were then conducted on CSD values. CSD values were decomposed into frequency-time domain using Fast Fourier Transform with a fixed window size of 1024 samples to generate a sequence of amplitude and phase components for the frequency bins between 6-12Hz. Due to the hypothesis of the present study, coherence analysis was restricted to the alpha range, defined here as 6-12Hz, a commonly used range in young children (McLaughlin, Fox, Zeanah, & Nelson, 2011; Oberman, Ramachandran, & Pineda, 2008).

Coherence was then computed in the form of event-related phase coherence (ERPCOH) from the aforementioned resting state epochs using the newcrossf function provided by EEGLAB (Delorme & Makeig, 2004). For each channel pair, phase coherence value was calculated by averaging ERPCOH of all latencies and of all the frequency bins encompassed by alpha band, resulting in 300 unique average alpha coherence values for every possible electrode pair.

### Statistical Analysis

In order to retain participants who only contributed EEG at one time point, analyses of EEG differences were conducted separately at 12 and 24 months of age.

#### Statistical Analysis: Group Comparisons

Two main group comparisons were performed: (1) TSC vs TD (2) ASD+ vs ASD−. As a reminder, all infants included in ASD group comparison had a TSC diagnosis. Identical analysis approaches (described below) were used for both of the group comparisons. Due to the large number of comparisons, type I errors were controlled using false discovery rate (FDR) methods (Benjamini & Hochberg, 1995). While uncorrected P values are reported throughout the results section for ease of interpretation, results are only determined to be (and described as) significant if they remained so following FDR correction (FDR=0.1). For any significant group contrasts, we confirmed that the relevant measurement values did not vary across acquisition sites (P<0.05).

Power: Instead of selecting predefined regions of interest, we took an unbiased, data-driven approach to examine power across the entire scalp. Group differences (in all 25 electrodes), were examined using test statistics which represented normalized differences in group means. A permutation test, where group labels were randomly assigned to subjects in each resample, was used to estimate the distribution of the test statistics. Two separate permutation tests (100,000 permutations each) were used to study group differences at 12 and 24 months.

PAF: Two separate repeated measure ANCOVA’s (with site entered as a covariate) were used to determine whether there were group differences in PAF at 12 and 24 months. To include all possible data and study regional differences, each region was investigated separately in follow-up analyses.

APC: We also applied non-parametric permutation testing to study group differences in APC. We examined APC for every possible electrode pair combination across the scalp (300 electrode pairs). Group differences in APC were examined using two separate permutation tests (described above) at 12 and 24 months.

#### Statistical Analysis: PAF and Cognitive Function

In order to maximize both sample size and range of cognitive function, correlations between cognitive function and PAF were collapsed across all infants (TD & TSC). A non-parametric Kendall’s tau correlation was employed to examine relationships between cognitive function and PAF for each separate region of interest (frontal, central & occipital), so as to make no assumptions about the distribution of the data.

## Results

### TSC/TD group comparison

Alpha Power: There were no alpha power differences between TSC and TD infants at 12 months (P>0.39) or 24 months (P>0.005).

PAF: At 12 months, there was no significant effect of group (F(1, 27)=2.322, P=.139), or region (F(2, 54)=.379, P=.69), and no group × region interaction effect (F(2, 54)=.257, P=.774) on PAF. At 24 months, there was a significant effect of group (F(1, 34)=5.909, P=.020), where PAF values were significantly lower in the TSC group compared to the TD group. There was also a significant effect of region (F(2, 68)=3.549, P=.034), but no group × region interaction effect (F(2, 68)=.042, P=.959). PAF values for each region of interest are described in Table 2.

**Table 2.** Peak alpha frequency values and group comparison results.

|  | TD | TSC | Test Statistic | p-value* |
| --- | --- | --- | --- | --- |
| **12-Months** |  |  |  |  |
| **Frontal** | 7.42 (0.53) n=18 | 7.16 (0.59)<br>n=16 | *U*=102 | .154 |
| **Central** | 7.49 (0.47) n=19 | 7.36(.48) n=16 | *U*=124.5 | .367 |
| **Occipital** | 7.25 (.58) n=18 | 6.9 (.63) n=16 | *U*=102.5 | .154 |
| **24-Months** |  |  |  |  |
| **Frontal** | 8.31 (.39) n=12 | 7.71 (0.71)<br>n=27 | *U*=75 | .007* |
| **Central** | 8.42 (.44) n=12 | 7.8 (0.87)<br>n=27 | *U*=86 | .026* |
| **Occipital** | 8.08 (.73) n=12 | 7.56 (0.91)<br>n=26 | *U*=112.5 | .174 |

\*Represents differences between TSC and TD that remained significant after FDR correction.

APC: At 12 months, infants with TSC (M=.24, SD=.03) showed significantly decreased APC compared to TD infants (M=.31, SD=.09) for a long-range temporal interhemispheric connection between electrodes T7 and C4 (P<0.00013; see Figure 1). There were no differences in APC for this connection between the two sites of acquisition (P>.232). At 24 months, there were no significant differences in APC (P>0.002).

**Figure 1.**
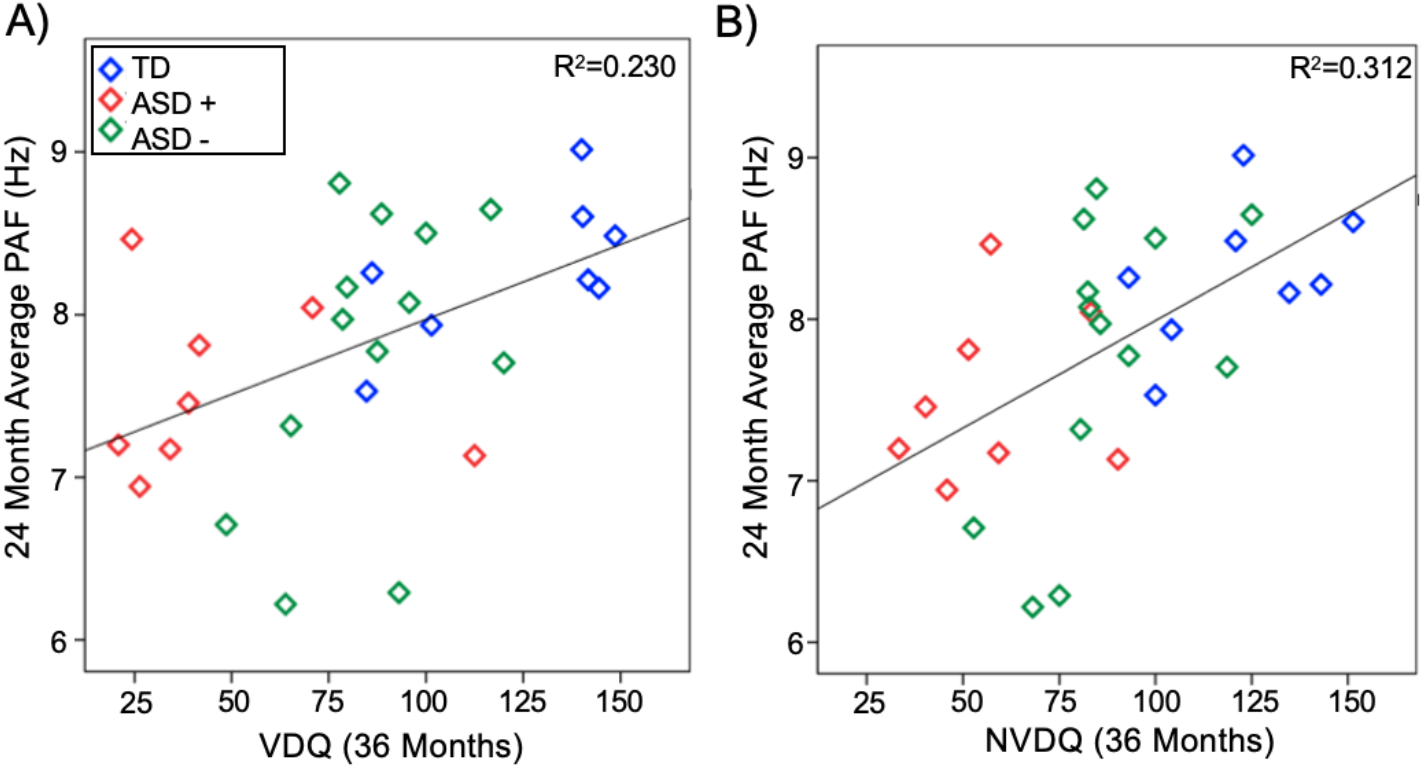
A) Log-transformed average power spectral density for both TSC (orange) and TD participants (blue). Shaded regions represent the median absolute deviation. B) Depiction of the interhemispheric connection that differentiated TD from TSC infants at 12 months of age (labelled connection represents significant electrode pair). C) Dot plot demonstrating APC for electrode pair T7-C4 (with group mean and error bars representing SD) for both TD and TSC groups.

### ASD+/ASD− group comparison

Alpha Power: There were no power differences between ASD+ and ASD− infants at any time point (12 months P>0.10; 24 months P>0.06).

PAF: At 12 months, there was no significant effect of group (F(1, 8)=1.172, P=.311), or region (F(2, 16)=.239, P=.790), and no group × region interaction effect (F(2, 54)=.822, P=.457). At 24 months, there was no significant effect of group (F(1, 21)=.248, P=.624). There was a significant effect of region (F(2, 42)=3.549, P=.026), but no group × region interaction effect (F(2, 42)=1.787, P=.180).

APC: There were no significant differences in APC between ASD+ and ASD− groups at 12 months (P>0.003). Decreased long range interhemispheric APC (between electrodes T9 and C4) was found in ASD+ (M=0.23, SD=0.03), compared to ASD− infants at 24 months (M=0.28, SD=0.04; P=0.00045; See Figure 2). Interestingly, this is a highly similar connection to that which differentiated TSC and TD participants at 12 months.

**Figure 2.**
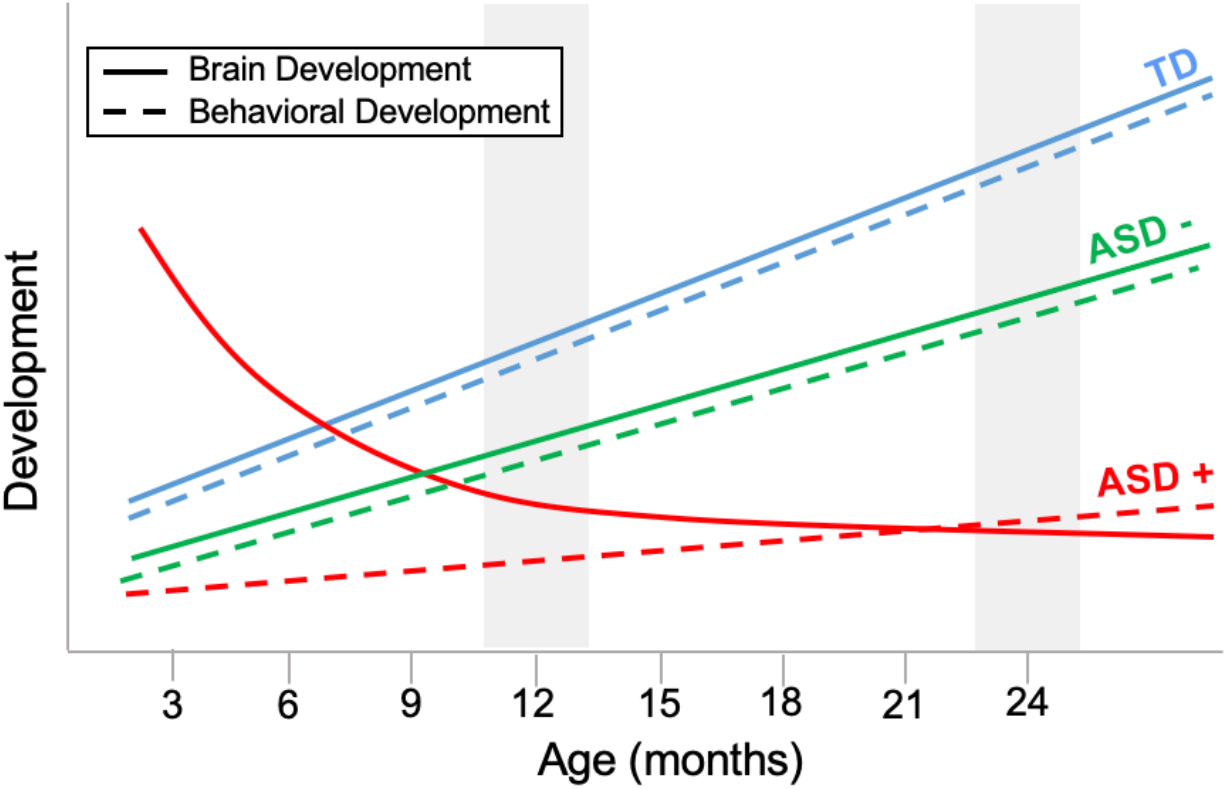
A) Log-transformed average power spectral density for both ASD+ (red) and ASD− participants (green). Shaded regions represent the median absolute deviation. B) Depiction of the interhemispheric connection that differentiated ASD+ from ASD− infants at 24 months of age (labelled connection represents significant electrode pair). C) Dot plot demonstrating APC for electrode pair T9-C4 (with group mean and error bars representing SD) for both ASD+ and ASD− groups.

### PAF & Cognitive Function

Kendall’s Tau correlations were performed to test the relationships between PAF (at 12 and 24 months), and cognitive function at 36 months across the entire sample (i.e., TD and TSC). 24 month PAF across all three regions of interest was found to be positively associated with NVDQ at 36 months, and frontal and central PAF at 24 months were found to be positively associated with VDQ at 36 months (see Table 2 & Figure 3).

**Table 3.** Associations between peak alpha frequency cognitive function across all participants.

|  | **VDQ** | **NVDQ** |
| --- | --- | --- |
| **12 months** | *t (P)* | *t (P)* |
| **Frontal** | .013 (.940) | -.026 (.879) |
| **Central** | .141 (.401) | .211 (.208) |
| **Occipital** | .199 (.234) | .258 (.123) |
| **24 months** |  |  |
| **Frontal** | .334 (.013)* | .420 (.002)* |
| **Central** | .426 (<.001)* | .459 (.001)* |
| **Occipital** | .236 (.079) | .314 (.020)* |

\*Represents associations that remained significant after FDR correction.

**Figure 3.**
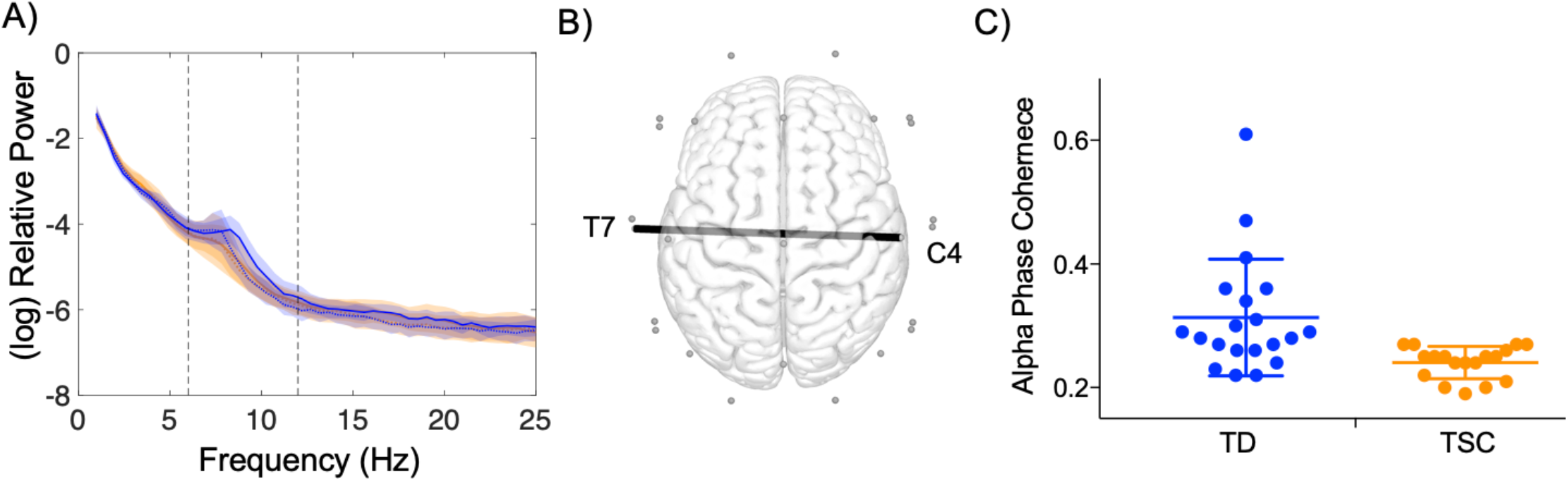
Scatter plots demonstrating the relationship between 24 month average PAF (averaged over frontal; central and occipital regions) and 36 months (A) VDQ, and (B) NVDQ.

## Discussion

Many infants with TSC show developmental delays and emerging signs of ASD by 12 months of age, with the most robust differences occurring between 12-24 months of age (Capal et al., 2017; Jeste et al., 2014; McDonald et al., 2017). Recent evidence from older children and adults with TSC suggests that widespread alterations in neuronal connectivity may underlie the emergence of neurodevelopmental disorders in TSC (Im et al., 2015; Lewis et al., 2012; Peters et al., 2012, 2013). The present study employed EEG measures of functional connectivity (PAF & APC) to investigate whether network disruption is present early in life in TSC, and whether PAF and APC differentiated TSC infants who did, and did not, later receive an ASD diagnosis. We also explored whether early PAF related to later cognitive function. We expected to identify reductions in both PAF and APC during the first two years in infants with TSC. Based on previous literature describing the relationship between PAF and cognition in ASD, we also expected to find a positive association between PAF and later cognitive outcomes (Dickinson et al., 2017). This was a clinically representative cohort, with a high incidence of epilepsy and associated medications as well as cortical tuber burden, thus reinforcing the value of these EEG biomarkers even in the context of these clinical comorbidities.

### TSC/TD Group Comparison

Using an unbiased, data-driven statistical approach, we found that infants with TSC demonstrate reduced interhemispheric APC between the left and right temporal regions compared to TD infants as early as 12 months of age. Notably, these differences were identified despite the absence of group differences in alpha power, suggesting that phase coherence provides increased sensitivity to functional disruptions at this early time point. In addition to APC differences, we demonstrate that PAF is consistently decreased in TSC, showing significant differences in frontal and central regions at 24 months. To the extent that APC and PAF index the development of neural networks (Wolfgang Klimesch et al., 2007; Valdés-Hernández et al., 2010), these findings suggest that functional networks show disruption from a young age in TSC.

The present findings specifically implicate aberrant long-range functional connectivity in TSC. One of the processes crucial to the integrity of large scale functional networks is the myelination of long range connections, which facilitates the efficient transfer of information over distributed brain regions (Fornari, Knyazeva, Meuli, & Maeder, 2007). The integrity of long range structural connections is reflected in measurements of functional network properties, with oscillations in the alpha range found to be particularly reflective of changes in white matter structure (Jann et al., 2012; Picci, Gotts, & Scherf, 2016; Teipel et al., 2009; Valdés-Hernández et al., 2010). Conversely, disorders of white matter are associated with disruptions in alpha oscillations, with decreased PAF demonstrated in vascular dementia (Moretti et al., 2004), multiple sclerosis (Leocani et al., 2000; Tewarie et al., 2014; Van der Meer et al., 2013), and white (but not gray) matter tumors (Gloor, Kalabay, & Giard, 1968; Goldensohn, 1979). Decreased PAF and reduced APC may therefore indicate delayed or atypical maturation of white matter during the infancy period, which is consistent with the disrupted interhemispheric white matter connections that have been identified later in TSC (Baumer et al., 2017; Im et al., 2015; Krishnan et al., 2010; Peters et al., 2012).

### ASD+/ASD− Group Comparison

A strikingly similar pattern (to that differentiating TSC/TD at 12 months of age) of decreased interhemispheric connectivity emerged at 24 months in ASD+, compared to ASD− infants. This pattern is consistent with the fact that ASD is largely associated with long range decreased connectivity during childhood and adulthood (Wass, 2011), and closely resembles connectivity disturbances observed in idiopathic ASD using identical recording procedures and analyses (Dickinson et al., 2018).

The timing of differences in connectivity patterns between ASD+ and ASD− infants highlighted by the present study indicates that these changes in connectivity are not static, and likely change over development, as we would expect given the dynamic interplay of genes, environment, and clinical comorbidities that affect neural function in these infants. In fact, in other high risk infant populations, it is the change in neural connectivity over the first year of life that distinguishes infants who develop ASD from those that do not (Shen & Piven, 2017). Moreover, in nonsyndromic ASD, imaging studies have demonstrated a pattern of under-connectivity during childhood that seems to be preceded by structural over-connectivity (Wolff et al., 2012) and early cortical overgrowth (Courchesne, 2004; Courchesne & Pierce, 2005; Hazlett et al., 2017), as well as increased functional connectivity (quantified using alpha coherence)(Wolff et al., 2012), in the first months of life.

Therefore, it is possible that the lack of significant connectivity differences at 12 months may reflect a transitional period in the infants who go on to develop ASD (see Figure 4). In nonsyndromic ASD, several neurobiological mechanisms have been attributed to underlie the atypical ‘crossover’ trajectory captured in ASD, including differences in axonal pruning and myelination (Courchesne, 2004; Courchesne & Pierce, 2005; Hazlett et al., 2017), differences in large scale anatomy (Orekhova et al., 2014), and a complex interplay between gray and white matter changes (Wolff et al., 2015, 2012). It should be noted that this is the first study to examine such change in neural function in TSC, and it supports the need for longitudinal investigation in neurodevelopmental disorders, regardless of underlying genetic etiology. Even though the impact of the TSC mutation impacts brain structure and function before birth, there are ongoing changes in the functional brain network that may be able to be modulated by early interventions (both pharmacological and behavioral). For instance, pharmacological intervention using an mTOR inhibitor, everolimus, appears to improve the microstructural integrity of the corpus callosum in patients with TSC (Courchesne, 2004; Hazlett et al., 2017).

**Figure 4.**
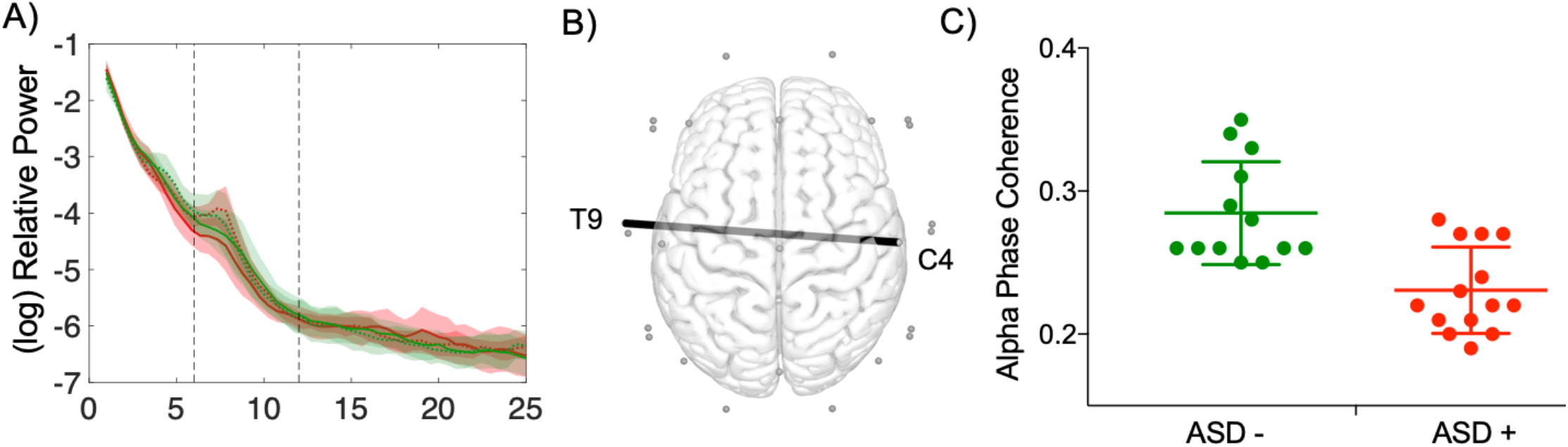
Theoretical (simplified) trajectories of brain and behavioral development across the first two years of life, with gray patches indicating timing of present study. Crucially, the hypothetical trajectories suggested here would elucidate why behavioral measures may discriminate ASD and no ASD in TSC at 12 months of age, but EEG measures do not.

### Cognition & PAF

Finally, we asked whether early measures of network function (PAF) related to individual differences in later clinically meaningful dimensional outcomes, which we quantified using measures of cognitive function. We found that across the entire sample, 24 month PAF was associated with cognitive functioning measured at a later timepoint (36 months). This is consistent with cross sectional work demonstrating that PAF is related to cognitive function in children with ASD (Bos et al., 2015; Ecker et al., 2016), and neurotypical adults (Peters et al., 2019; Tillema, Leach, Krueger, & Franz, 2012), and extends these findings to demonstrate that PAF also holds value as an early prognostic marker related to later developmental outcomes. While the limited sample size in the present study precluded within-subgroup analyses, future studies will explore whether PAF is able to predict cognitive outcomes within (as well as across) clinical subgroups.

### Limitations

Structural lesions such as cortical and subcortical hamartomas (tubers) may alter EEG measures of functional connectivity through volume conduction (Dickinson et al., 2017). We applied surface Laplacian transform in an attempt to mitigate the effects of tubers (Grandy et al., 2013; W. Klimesch, Schimke, & Pfurtscheller, 1993; Richard Clark et al., 2004). The surface Laplacian isolates the source activity under each electrode. Therefore, current source density (CSD) estimates capture the unique properties of each electrode while minimizing activity that is broadly distributed across multiple electrodes thereby reducing individual differences in volume conduction introduced through cortical differences, including tuber burden. The special consideration given to this specific signal processing issue resulted in the detection of individual differences in subgroups of TSC infants (ASD+/ASD−), two groups equivalently impacted by tuber burden.

The presence of interictal EEG abnormalities in infants with epilepsy also can alter the EEG signal being measured (Nunez & Srinivasan, 2006; Nunez et al., 1997). However, the presence of seizures is more often associated with EEG hypersynchrony (Kayser & Tenke, 2015), which would manifest as increased phase coherence. Therefore, the fact that TSC participants in the present study show decreased phase coherence would suggest that these differences are not spuriously introduced through the presence of seizures. In addition, our goal was to include a clinically representative sample of infants with TSC, and the findings reinforce the fact that EEG biomarkers in early infancy, as in other high risk populations, hold tremendous value for early risk stratification, prediction and as possible biomarkers of change with intervention.

### Future Directions: APC and PAF as markers of altered neural connectivity in TSC

The present results indicate that both APC and PAF hold utility in early life in TSC. Both of these metrics not only are able to differentiate TSC from TD infants, but are also sensitive to differences in infants who later develop ASD, and show early differences according to later cognitive function. APC and PAF have been previously associated with measures of structural network development, particularly white matter connections. In addition, a large literature of child and adult studies in TSC find disruptions in white matter integrity. Considering both of these bodies of literature, altered PAF and APC in the present study may be capturing early divergences in white matter development.

The demonstration that functional connectivity deficits in TSC can be measured as early as 12 months and relate to neurodevelopmental outcomes suggests that EEG could serve as a robust tool not only to predict seizures (which is already being done clinically) but also to inform our understanding of neurodevelopmental disorders in TSC. Future research is needed to examine functional EEG measures alongside white matter maturation in TSC in infancy in order to better understand the neurobiological mechanisms underlying early network disruption. Studying infants from earlier in life will also allow us to determine whether TSC ASD+ infants show similar patterns of long range hyperconnectivity to those seen during the first months of life in infants at familial risk for ASD. The in-utero diagnosis of TSC certainly facilitates this research avenue. However, due to the rare nature of TSC, and further reduction of sample sizes that occurs when comparing TSC infants based on ASD outcomes, sufficiently large samples will only be achieved through multi-site studies.

### Conclusions

Here we find that EEG measures of brain network function are altered in TSC and those infants with TSC who develop ASD in the first two years of life. Specifically, we find 1) decreased long range coherence in TSC at 12 months and lower peak alpha frequency at 24 months, 2) decreased long range coherence at 12 months in infants who develop ASD, and 3) an association between peak alpha frequency and cognitive function at age 3. The identification of objective, quantifiable markers of atypical development within the first years of life in TSC can improve the timing and accuracy of identification of those infants requiring additional, targeted developmental interventions from very early in life. The availability of scalable objective markers of altered neural development is particularly advantageous during infancy, as this is a period of peak neural plasticity during which developmental trajectories could potentially be steered. Thus, while a formidable challenge, particularly in the context of a rare disorder marked by high heterogeneity, this early detection approach is highly warranted considering the potential clinical and translational applications associated with the identification of objective (and potentially, prognostic), biological markers of neurodevelopmental outcomes.

## Acknowledgements

The authors wish to thank all of the infants and families who generously participated in the study. The present research was supported by grants from the Department of Defense (DOD CDMRP TSCRP: 2011–2014) and the UCLA CTRC (UL1TR000124). The authors are also grateful to Scott Huberty and Lauren Baczewski for their role in the study, and to Manjari Daniel for her helpful comments on the manuscript.

## Conflict of Interest

S.S.J. serves as a consultant for Roche Pharmaceuticals and on the professional advisory board for the Tuberous Sclerosis Alliance. M.S. receives research support from Novartis, Roche, Pfizer, LAM Therapeutics and Quadrant Biosciences, and has served on the Scientific Advisory Board of Sage Therapeutics, Roche, Celegne and Takeda. The remaining authors (A.D, K.J.V., C.A.N.) do not have any conflicts of interest associated with this study.

## Notes

#### Summary of Updates

Missing reference.

