## Supplementary Information for "Early patterns of functional brain development associated with autism spectrum disorder in tuberous sclerosis complex" (from another version of this preprint: version 1, 2019-03-16)

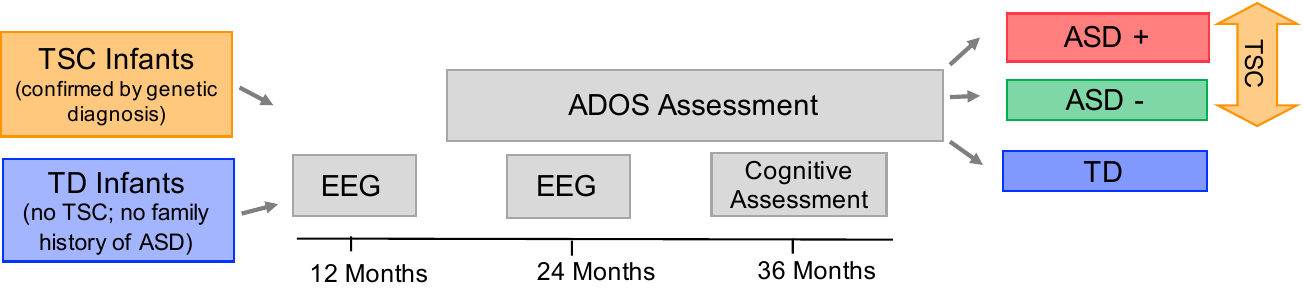

*Supplementary Figure 1.* A schematic diagram of the current study protocol.

Supplementary Table 1. *Seizure and medication information for infants with TSC.*

|  | **TSC** | **ASD+** | **ASD-** | **Test Statistic*** | **p-value*** |
| --- | --- | --- | --- | --- | --- |
| Age at seizure onset in months,  Median [Interquartile range] (n) | 4.00 [4.56]  (29/32) | 4.00 [5.77]  (15) | 4.00 [3.81]  (13) | *U*=107.5 | .650 |
| History of infantile spasms (n) | 62.5%  (20/32) | 62.5%  (10/16) | 60%  (9/15) | χ^2^=.02 (1) | .886 |
| **12-Months** | | | | | |
| Median number of AEDs [Range] (n) | 2 [0-4]  (22) | 2 [0-4]  (13) | 1.50 [0-2] (8) | *U*=39.00 | .374 |
| Past or current seizures (n) | Current: 58% (14)  Past: 17% (4)  Never: 25% (6) | Current: 62% (8)  Past: 15% (2)  Never: 23% (3) | Current: 58% (7)  Past: 17% (2)  Never: 25% (3) | χ^2^=.034 (2) | .983 |
| **24-Months** | | | | | |
| Median number of AEDs [Range] (n) | 2 [0-4]  (26) | 2 [0-4]  (13) | 2 [0-4]  (13) | *U*=72.50 | .545 |
| Past or current seizures (n) | Current: 58% (15)  Past: 34% (9)  Never: 8% (2) | Current: 43% (6)  Past: 57% (8)  Never: 0% (0) | Current: 64% (9)  Past: 22% (3)  Never: 14% (2) | χ^2^=3.25 (2) | .197 |

*Represents the comparison between TSC/ASD and TSC/no ASD.

Supplementary Table 2. *Additional seizure and medication information for infants with TSC at each age that EEG data were acquired.*

|  | **TSC** | **ASD+** | **ASD-** | **Test Statistic*** | **p-value*** |
| --- | --- | --- | --- | --- | --- |
| **12-Months** | | | | | |
| Medication type | VGB (12), LEV (8), OXC (6), TPM (4), CBZ (2), VPA (2), CLD (1), GBP (1), LMT (1), PHB (1), None (4),  Missing (17) | LEV (8), VGB (7), OXC (3), TPM (3), CBZ (2), LMT (1), CLD (1), GBP (1), None (2),  Missing (5) | VGB (5), VPA (2), OXC (3), PHB (1), TPM (1),  None (2),  Missing (12) | - | - |
| Frequency of seizures | Daily: 40% (8)  Weekly: 0% (0)  Monthly: 15% (3)  Never: 45% (9) | Daily: 36% (4)  Weekly: 0% (0)  Monthly: 18% (2)  Never: 46% (5) | Daily: 50% (5)  Weekly: 0% (0)  Monthly: 10% (1)  Never: 40% (4) | χ^2^=.536 (2) | .765 |
| Seizure status (in response to medication) | Full: 19% (3)  Partial: 31% (5)  Refractory: 50% (8) | Full: 20% (2)  Partial: 40% (4)  Refractory: 40% (4) | Full: 14% (1)  Partial: 14% (1)  Refractory: 72% (5) | χ^2^=1.75 (2) | .418 |
| **24-Months** | | | | | |
| Medication type | EVL (1), VGB (16), LEV (10), OXC (10), CBZ (5), TPM (3), VPA (2), RFM (1), SRL (1), PHT (1), TAU (1), ZNS (1), None (4), Missing (12) | VGB (10), LEV (7), OXC (5), CBZ (3), TPM (2), TAU (1), ZNS (1),  None (1)  Missing (4) | VGB (6), OXC (5), LEV (2), CBZ (2), VPA (2), RFM (1), TPM (1), SRL (1), EVL (1), PHT (1), None (3),  Missing (8) | - | - |
| Frequency of seizures | Daily: 26% (6)  Weekly: 22% (5)  Monthly: 4% (1)  Never: 48% (11) | Daily: 8% (1)  Weekly: 23% (3)  Monthly: 8% (1)  Never: 61% (8) | Daily: 42% (5)  Weekly: 16% (2)  Monthly: 0% (0)  Never: 42% (5) | χ^2^=3.92 (3) | .270 |
| Seizure status (in response to medication) | Full: 35% (8)  Partial: 22% (5)  Refractory: 43% (10) | Full: 50% (7)  Partial: 29% (4)  Refractory: 21% (3) | Full: 27% (3)  Partial: 9% (1)  Refractory: 64% (7) | χ^2^=3.86 (2) | .145 |

*Represents the comparison between ASD+ and ASD-.

Abbreviations: TSC = Tuberous sclerosis complex; ASD = Autism spectrum disorder; ACTH = Adrenocorticotropic hormone; CLD = Clonidine; CBZ = Clobazam; CLZ = Clonazepam; EVL = Everolimus; GBP = Gabapentin; LEV = Levetiracetam; LMT = Lamotrigine; OXC = Oxcarbazepine; PHB = Phenobarbital; PHT = Phenytoin; RFM = Rufinamide; SRL = Sirolimus; TAU = Taurine; TPM = Topiramate; VPA = Valproic Acid; VBG = Vigabatrin; ZNS = Zonisamide;
